# High-resolution 7T fMRI reveals the visual sensory zone of the human claustrum

**DOI:** 10.1101/2023.09.18.558213

**Authors:** Adam Coates, David Linhardt, Christian Windischberger, Anja Ischebeck, Natalia Zaretskaya

## Abstract

The claustrum is a thin subcortical gray matter structure located between the insula and the putamen. It has numerous bilateral connections with the cortex and is thought to play an important role in higher-level aspects of perception and cognition, with hypotheses including multisensory integration, attention and consciousness. The claustrum’s thin shape makes it difficult to investigate, leaving the hypothesis regarding its function largely untested. In the current study, we used high-resolution ultra-high field (7 Tesla) functional magnetic resonance imaging (fMRI) to measure claustrum activity in human participants, while they were presented with visual, auditory or audiovisual naturalistic stimuli. We found distinct visual responses in the claustrum at a spatial location that was consistent across participants, hemispheres and scanning sessions. This is the first study to demonstrate evoked sensory responses within the human claustrum. It opens the possibility for future noninvasive investigation of the claustrum’s role in sensory processing.

## Introduction

The claustrum is a thin bilateral subcortical structure that is situated between the putamen and the insula and is interconnected with most cortical and subcortical areas within the mammalian brain ^1–3^. Human in vivo diffusion tensor imaging (DTI) revealed that the claustrum is the most connected region of the brain per volume unit ^4^. The claustrum has also been shown to have a high density of serotonin 2a receptors (5-HT_2A_), the primary targets of psychedelic drugs ^5,6^. As a result, administering psilocybin leads to widespread changes in resting-state functional connectivity of the claustrum with the rest of the brain ^7^. While much attention has been given to the structural properties of the claustrum and its connectivity with the rest of the brain, the function of the claustrum remains elusive.

The unique structural properties of the claustrum and its connections have led to a prominent hypothesis about its role in multisensory integration, binding and consciousness (Crick & Koch, 2005). According to Crick and Koch, the unique sensory-to-claustrum and subsequent claustrum-to-frontal type connections are critical in coordinating the signals from sensory cortices that are ultimately fed forward to higher brain areas such as the prefrontal and frontal cortex. However, the empirical support for this hypothesis remains weak. A single-cell electrophysiology study examining responses of the claustrum neurons to visual and auditory stimuli in the macaque failed to find any signatures of multisensory integration ^9^. A single case study of a patient suffering from epilepsy reported that electrical stimulation close to the claustrum resulted in temporary non-responsiveness and loss of consciousness ^10^. However, a replication with five subjects with electrodes within the claustrum found no effect on consciousness ^11^. Despite these mixed findings, disruption to sensory processing as a result of claustrum lesions or electrical stimulation has been a prominent observation across the literature ^12^.

Research on the rodent claustrum primarily points to its role in attention. For example, it was shown that optogenetic inhibition of claustrum neurons in the mouse did not affect task performance in a task without a distractor stimulus^13^. However, task performance decreased when an auditory distractor tone was presented during claustrum inhibition. An additional recording of activity in the auditory cortex showed that optogenetic stimulation of particular claustrum cells led to auditory cortex suppression in response to the distractor tones. The authors concluded that the claustrum inhibits irrelevant sensory stimuli that are not required to perform a task, making an individual more resilient to distraction ^13,14^. Given the differences in overall brain anatomy and cognitive function between humans and rodents ^15,16^, it still remains unclear to which extent such findings are generalizable to humans.

A well-documented property of the claustrum’s functional organization is the existence of sensory zones ^9,17^ with corresponding topographically organized claustrocortical connections ^18–20^. Specifically, auditory, visual and somatosensory zones have been described in the claustrum, with neurons preferring sensory input of the corresponding modality. The claustrum’s thin shape and its location deep within the brain are challenging for conventional neuroimaging in human participants.

In this study, we utilized ultra-high field 7T fMRI to determine whether it is possible to elicit measurable visual and auditory responses in the human claustrum. We presented naturalistic visual, auditory and audiovisual stimuli to participants in a 7T fMRI experiment. We expected to find modality-specific zones within the claustrum that responded to either visual or auditory stimulation and potentially also signatures of multisensory integration, which would show up as response enhancement in multisensory compared to unisensory conditions.

## Results

Our aim in the current study was to determine whether high-resolution 7T fMRI would allow us to identify visual and auditory sensory zones of the human claustrum, and whether these or any other regions of the claustrum show signs of audio-visual integration. To test this, we presented participants with naturalistic video clips containing visual only, auditory only or audiovisual information (see **Fig.1a** for an example), while measuring fMRI activity in either the left or the right claustrum. To unambiguously assign functional activations to the claustrum, we manually labelled the left and right claustrum in an ultrahigh-resolution post-mortem MRI image that was mapped to MNI space (see **Fig.1b** and Methods for details).

### Unisensory responses

We performed a group analysis across all voxels within the claustrum for the left and right claustrum separately, comparing responses of each unisensory condition with baseline. Our analysis revealed 2 significant clusters of voxels that responded stronger to visual stimulation compared to baseline in the left claustrum (most significant voxel in cluster 1; z-statistic = 3.3, p<0.001, d = 1.7, MNI x= -32.2, y= -3.75, z= -12, cluster size = 132.9 mm^3^, most significant voxel, cluster 2; z-statistic = 3.2, p<0.01, d = 1.6, MNI x= -37.5, y= -1.5, z= -18.8, cluster size = 21.9 mm^3^). We also found 1 significant cluster in the right claustrum that responded stronger to visual stimulation compared to baseline (most significant voxel in cluster; z-statistic = 3.9, p<0.001, d = 4.4, MNI x= 32.2, y= -5.25, z= -11.2, cluster size = 51 mm^3^) (**Fig. 1c**).

**Fig. 1.**
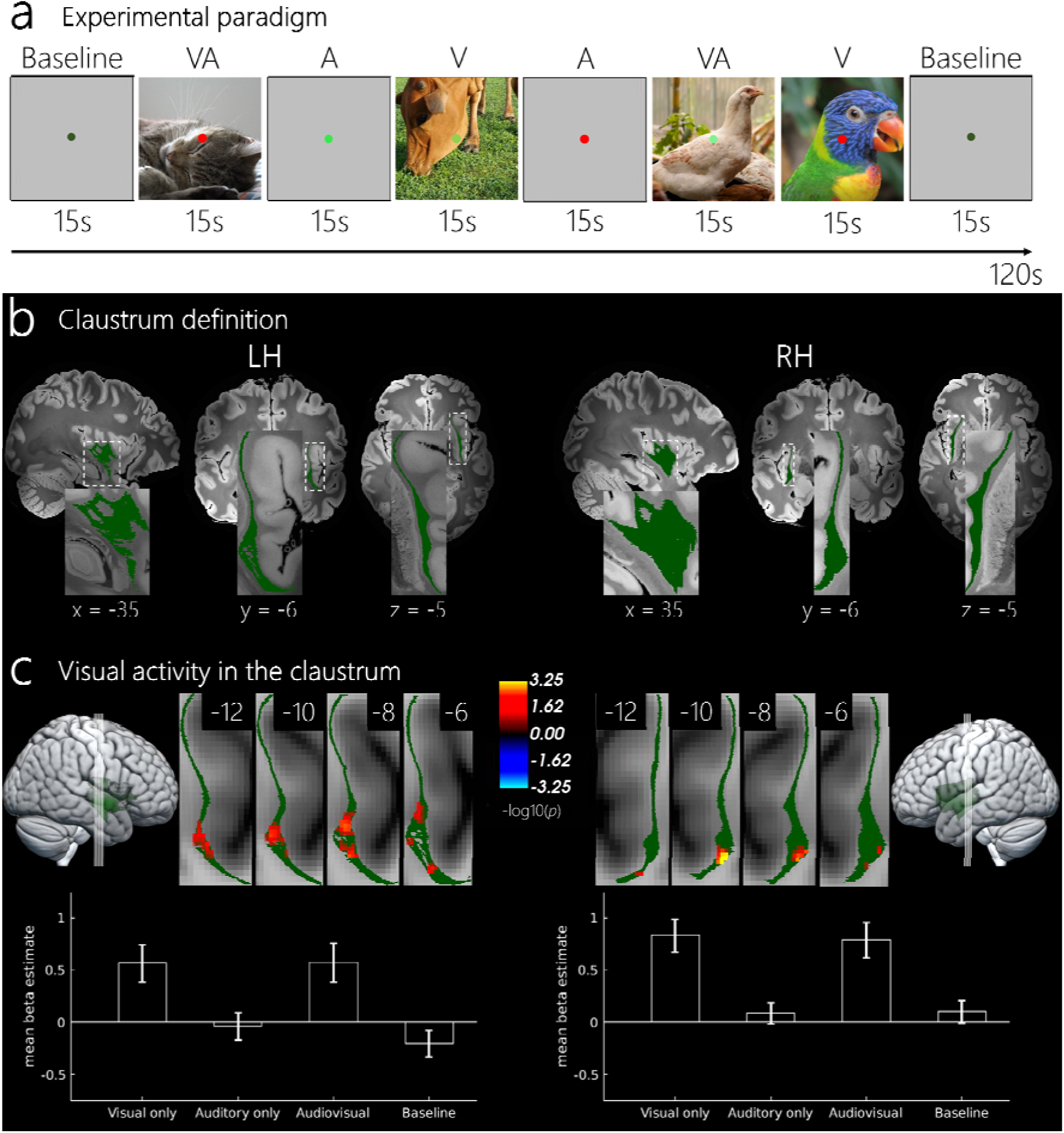
*a) An example of a typical condition sequence within a run. Each condition lasted for 15 seconds. The alternation of 1 baseline followed by 6 stimulus trials was repeated 4 times in each of the 6 runs for each session. Participants were instucted to fixate on the cental fixation point and to respond when the point changed to the color red. b) Example of the right and left claustrum label overlayed with the ultrahigh-resolution MRI image, which was used to identify the claustrum. We manually labelled the left and right claustrum twice and the union of the 2 claustrum labels per hemisphere is shown in green. Coordinates are shown in MNI space. c) Visually evoked activity for the left and right claustrum. Four coronal slices represent the size and location of the clusters which are corrected for multiple comparisons at a cluster forming threshold of p<0.01 and a cluster-wise p-value of p<0.05. Coordinates are shown in MNI space. Bar plots represent the mean beta estimates for each condition within the significant voxels. Error bars represent standard error of the mean (SEM).*

### Individual-level results

We then ensured that significant visual activity observed in the claustrum at the group level is present in each individual subject at a similar spatial location in unsmoothed data. To do this, we analyzed session 1 and session 2 of each subject together and then extracted MNI coordinates of the most significant voxel for the visual vs. baseline comparison. Peak coordinates of each subject as well as the mean and SD are shown in Table 1. Individual subject’s statistical maps are shown in a coronal view (left; y = -11.80, right; y = -6.55) in **Fig. 2**.

**Fig. 2.**
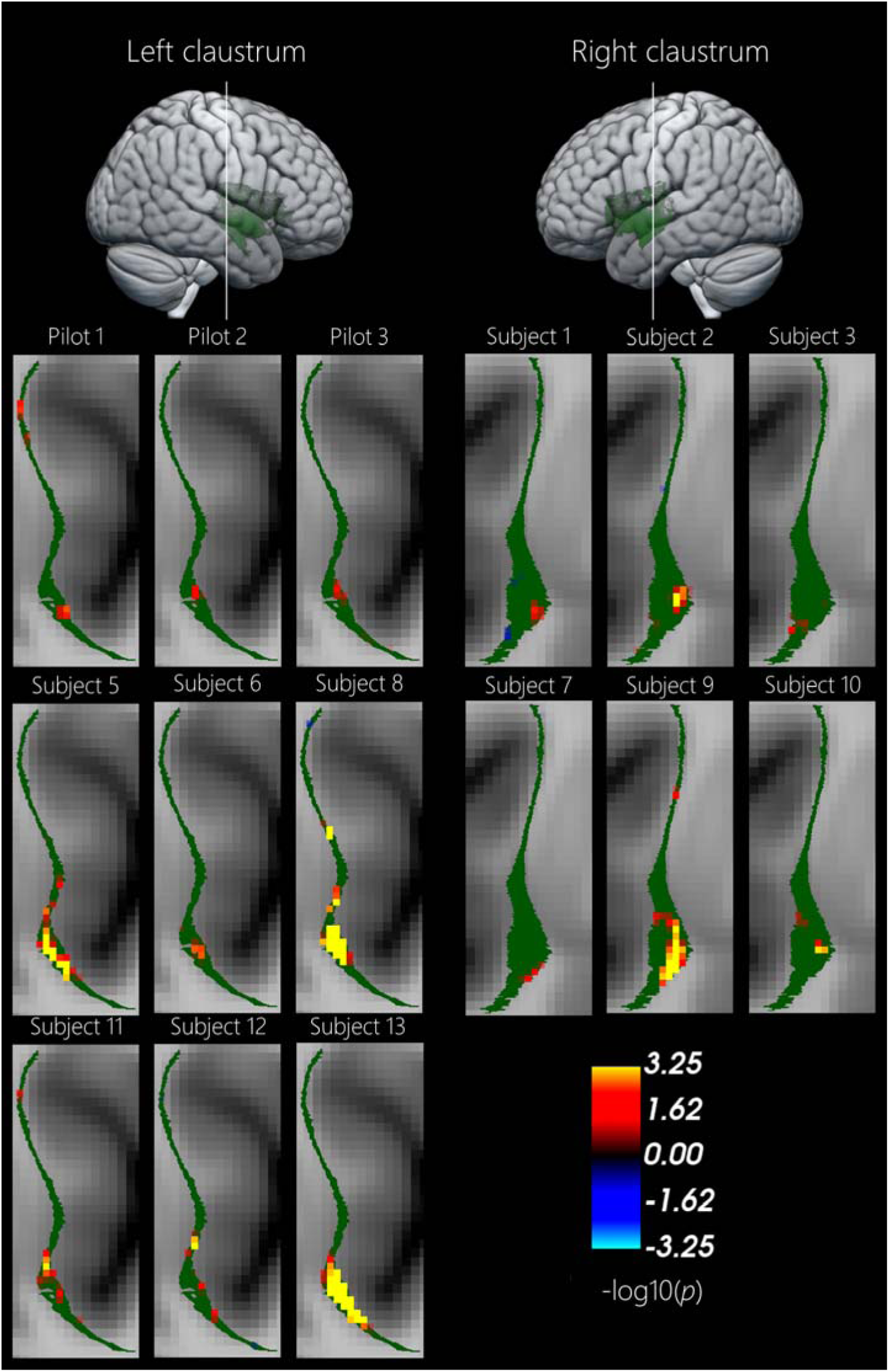
*Results of individual subjects in MNI space shown on an MNI template in coronal view. Left hemisphere response at y = -11.80 slice. Right hemisphere response at y = -6.55.*

**Table 1.**
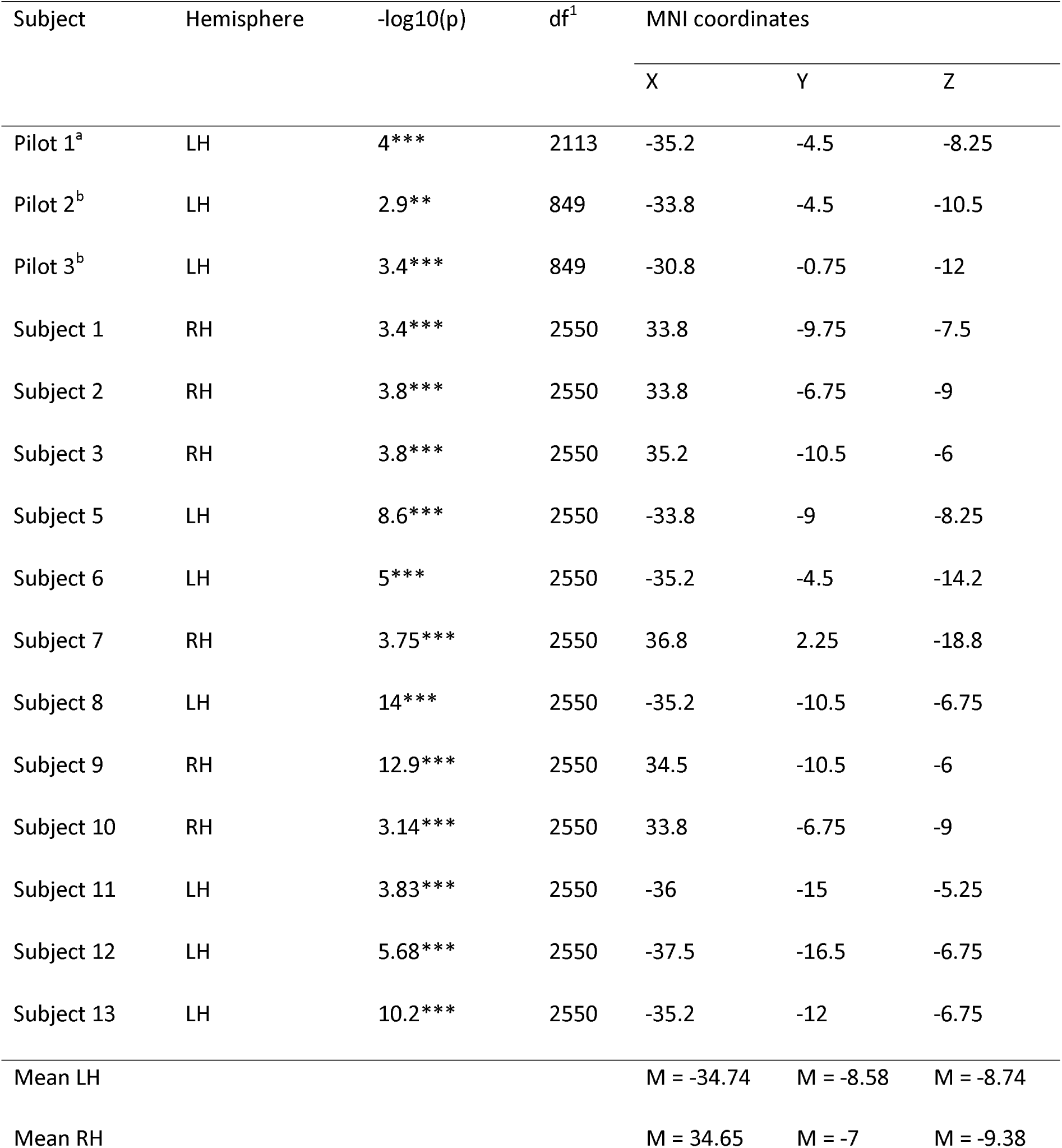
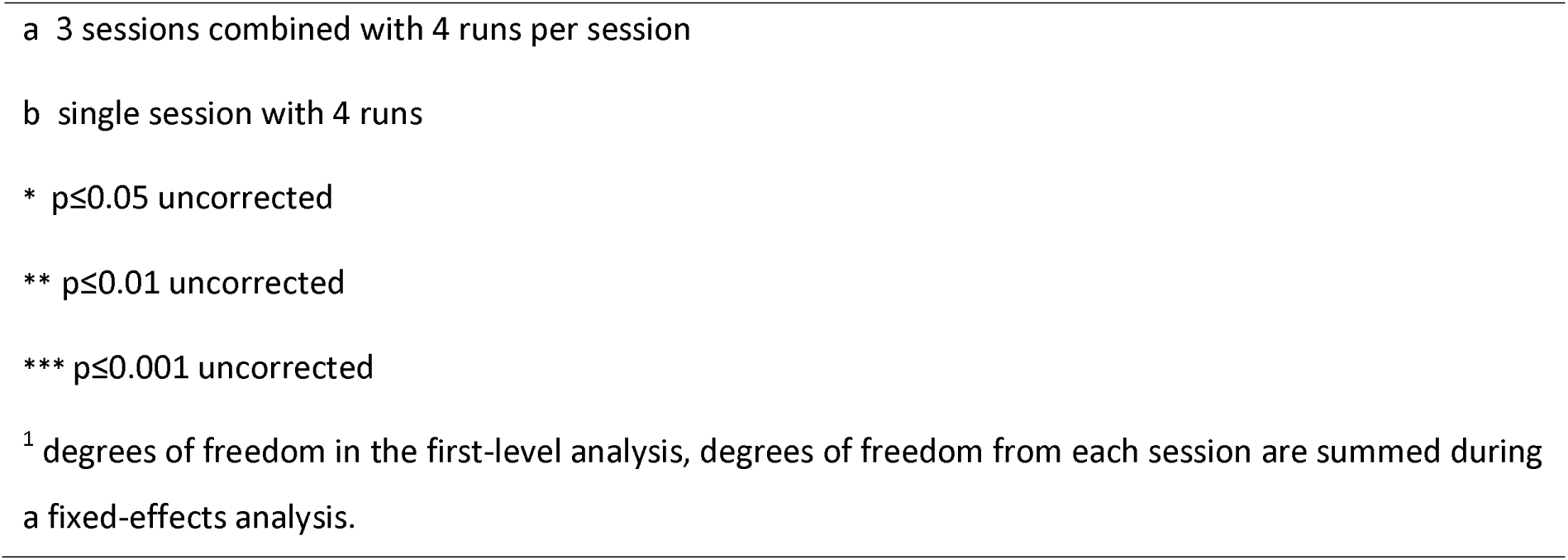
Visual activations in individual subjects.

### Auditory responses in the claustrum

We found no voxel clusters responsive to auditory condition after the multiple comparison correction in either hemisphere. Interestingly, we observed a significant cluster of voxels that showed a suppression of activity relative to baseline during the auditory stimulation, but only in the right hemisphere (most significant voxel in cluster: z-statistic = -3.4, p<0.001, d = 3.1, MNI x = 34.50, y = 0.75, z = -5.25, cluster size = 55.7 mm^3^, **Fig. 3a**). There were no voxels suppressed by the auditory stimulation in the left hemisphere even with a more liberal cluster-forming threshold of p<0.05. The auditory deactivations were not overlapping with visual activations (**Fig. 3b**).

**Fig. 3.**
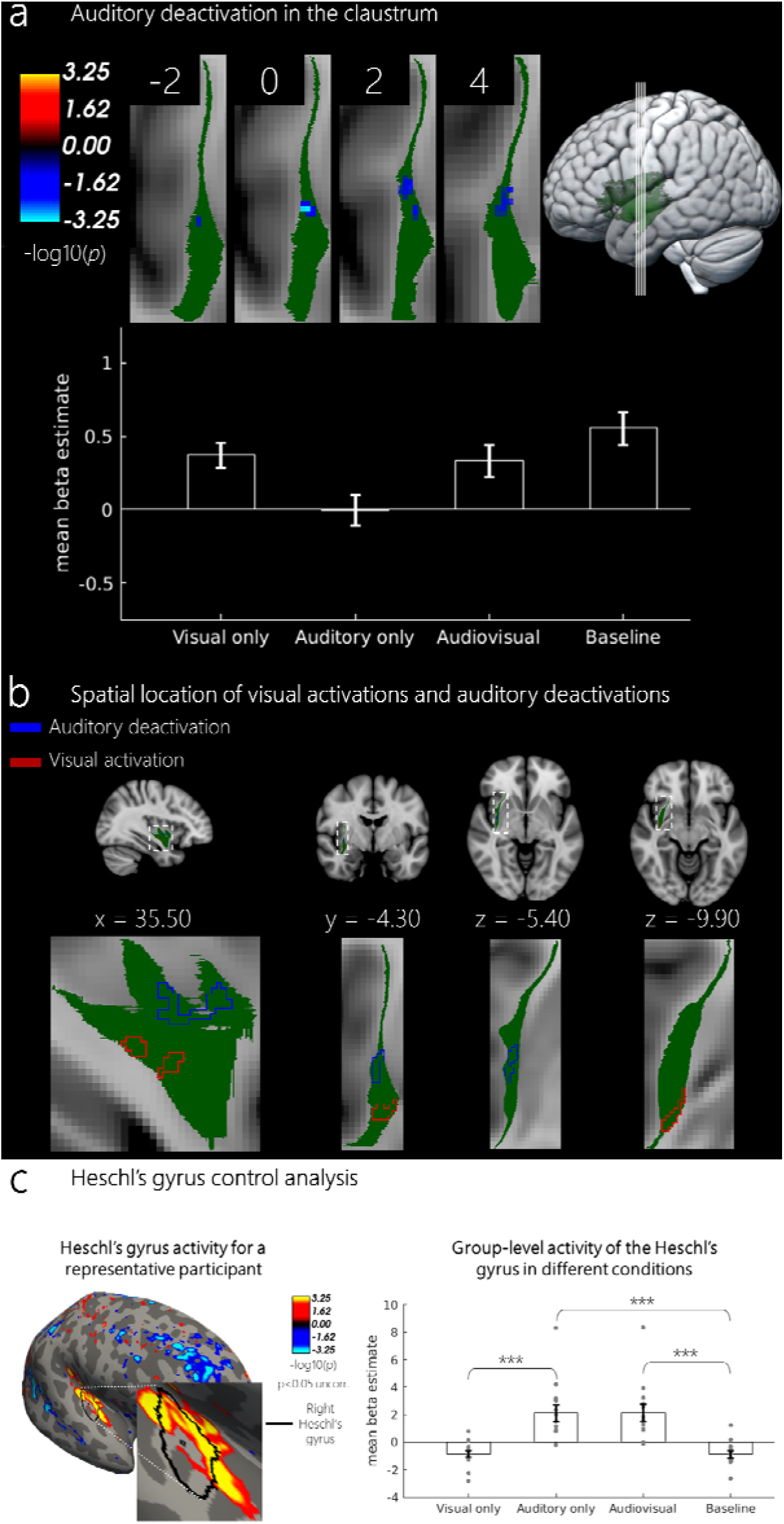
*a) Deactivations evoked by the auditory stimuli in the right claustrum. The reported cluster is corrected for multiple comparisons at a cluster forming threshold of p<0.01 and a cluster-wise p-value of p<0.05. Coordinates represent the center of mass for the cluster. Bar plot represents the mean beta estimates for each experimental condition within the significant voxels. Error bars represent SEM. b) Right hemisphere with the clusters for the visual activations and auditory deactivations shown as outlines to demonstrate that the clusters are spatially non-overlapping. Individual subject values represent an average activity of two sessions. Error bars represent SEM. ***p<0.001. c) An example of a single participant’s reconstructed surface (subject 2) showing significant auditory activation in the first session (p<0.05, uncorrected) in the right Heschl’s gyrus. Bar plot represents the group-level mean beta estimates for the Heschl’s gyrus for each experimental condition (N = 15).*

### Auditory cortex results

Since our main analysis did not reveal any significant activity in response to auditory stimulation that survived multiple comparisons correction, we ensured that our auditory stimuli were efficient in evoking auditory activity by analyzing responses of the auditory cortex. We extracted the mean beta estimates for each condition (visual, auditory, audiovisual and baseline) from the transverse temporal gyrus (Heschl’s gyrus), which corresponds to the primary auditory cortex. As expected, the primary auditory cortex exhibited no significant activation in response to visual stimuli (V-baseline, M = -0.84, SE = 0.25; t(14) = 0.24, p = 0.81, d = 0.02), but a significant activation in response to auditory stimuli (A-baseline, M = 2.13, SE = 0.62; t(14) = 5.4, p<0.001, d = 1.92), to audiovisual stimuli (AV-baseline, M = 2.17, SE = 0.6; t(14) = 5.50, p<0.001, d = 1.98) and when comparing auditory and visual stimuli (V-A, t(14) = 5.23, p<0.001). These results are summarized in **Fig. 3b**. It is therefore unlikely that the absence of auditory activity within the claustrum is related to the inefficient auditory stimulation in our experiment.

### Multisensory responses

We also looked at multisensory responses within the claustrum at the group level. We wanted to determine if there are claustrum regions beyond the unisensory zones that show multisensory responses. Although we found a cluster that showed an activity pattern consistent with the superadditive response ((AV + baseline) – (A + V), a closer examination of the cluster location and the beta estimates for each condition revealed that first, the cluster largely overlaps with the location of auditory deactivations (**Fig. 3a**) and second, the multisensory contrast effects are driven by suppression of auditory responses below the baseline (**suppl. Fig. 1**). The latter is inconsistent with a superadditive multisensory effect, which requires unisensory responses to be higher than the baseline ^22,23^. We therefore did not observe any signs of multisensory integration within the claustrum.

### Session-to-session consistency

Our main analysis thus reveals consistent bilateral visual responses at a specific location within the human claustrum. To check whether the location of visual activity observed at the group level was consistent within individual subjects across different scanning days, we performed a session-to-session consistency analysis of visual activity, additionally including auditory effects for completeness. We used one session to define a group of visual/auditory-selective voxels and measured the responses of these same voxels in the other session, statistically comparing the responses with zero.

We found that visual responses were consistent from session-to-session, as indicated by above-zero contrast estimates for both sessions (**Fig. 4a**). A one-sample t-test comparing the effects in each session against 0 revealed a significant effect for session 1 (t(11) = 4.56, p<0.001, d = 0.71) and a significant effect for session 2 (t(11) = 4.86, p<0.001, d = 0.59). In contrast, there was no significant auditory activity, neither for session 1 (M = 0.21, SE = 0.12; t(11) = 1.69, p = 0.12, d = 0.22) nor for session 2 (M = 0.17, SE = 0.16; t(11) = 1.06, p = 0.31, d = 0.16) which is consistent with the absence of auditory activations within the left and right claustrum at the group level (**Fig. 4b**). For the auditory deactivations Figure, we looked at the sessions-to-session consistency only within the right hemisphere that showed significant deactivation effects at the group level (**Fig. 4c**). However, we could not confirm the session-to-session consistency of these effects, as there was no significant difference from 0, neither for session 1 (M = -0.22, SE = 0.12; t(5) = -1.80, p = 0.13, d = 0.27), nor for session 2 (M = -0.24, SE = 0.14; t(5) = -1.78, p = 0.13, d = 0.29).

**Fig. 4.**
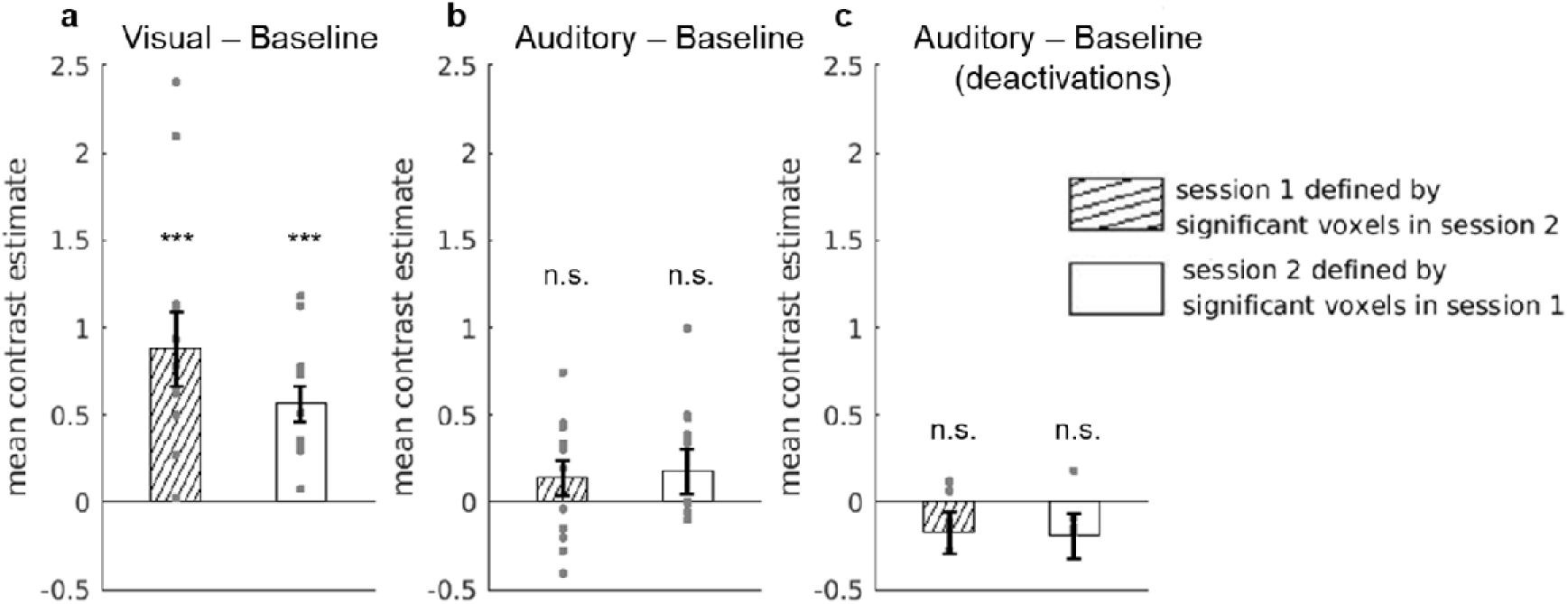
*Consistency of visual and auditory responses between session 1 and session 2. a) Visual activity for each session shows as the mean contrast estimates for the V – baseline comparison (activation). b) Auditory activity for each session shown as the mean contrast estimates for the A – baseline comparison (activation). c) Auditory suppression for each session shown as the mean contrast estimates for the A – baseline comparison (deactivation for the right hemisphere only with N = 6). Error bars represent standard error of the mean. (Note that pilot subjects were not included in this analysis, yielding (N = 12, for a & b). ***p<0.001.*

### Behavioral results

To ensure that participants were keeping their gaze on the fixation point throughout the experiment we calculated percentage of hits and reaction times. As expected, we found a high average percentage of hits (M = 86.95%, SD = 17.93%, IQR = 79.96% – 97%) across all participants and reaction times were less than 1 second (M = 0.67 s, SD = 0.15 s, IQR = 0.65 s – 0.70 s).

## Discussion

In this study, we aimed to investigate the visual, auditory and audiovisual sensory responses within the human claustrum using ultra-high resolution 7T fMRI and naturalistic video clips. We found visual responses within the claustrum that were consistent across sessions and appeared at a similar location across participants in both hemispheres. We did not find significant auditory activity and no response pattern consistent with audiovisual integration. These results provide the first insight into the modality-specific sensory responses within the human claustrum that have otherwise been demonstrated only in animal models.

Our findings of visually evoked activity within the claustrum are in line with what is known about claustrum physiology from animal models. For example, similar to a previous study that found visually responsive neurons at the more ventral claustrum locations ^9^, we also observed visual activity within a ventral claustrum site. Notably, a more ventral location of visual activity is also expected based on the topography of claustrocortical connections known from human tractography studies ^18,24^. While that study used natural movie clips, earlier studies using more controlled visual stimulation could also narrow down the feature-specific nature of the cells within the visual region of the cat claustrum ^25^. Neurons in the visual zone of the cat claustrum showed preference for elongated moving bar stimuli as opposed to stationary stimuli and were particularly selective for the orientation of the bar stimuli. These animal model studies suggest that neurons in the visual claustrum zone may show a preference for specific features of the visual stimuli. We intentionally used natural stimuli that contain a wide range of visual features. It therefore remains unknown if particular features of the stimuli used may have evoked a stronger response in the visual zone of the claustrum compared to other features. Future research will have to determine what stimuli features the visual zone is more selective to in humans.

In contrast to our expectations based on primate literature ^26^, we did not find any auditory activation within the claustrum in response to auditory stimulation. There may be different reasons as to why auditory responses could not be found. Firstly, it is likely that the auditory zone is located in a thinner part of the claustrum compared to the visual zone. The auditory zone identified described in primates, was located more dorsally compared to the visual region ^9^. Naturally, the claustrum’s structure becomes much thinner in the dorsal areas ^27^. The auditory zone is thus expected to be more susceptible to partial volume effects and to produce a weaker function signal, which we may have been unable to detect in the current experiment even using high spatial resolution at 7 Tesla. Future investigations that aim to differentiate between the auditory and the visual zone activity may require strategies that further increase the resolution of functional images ^28–31^.

Another potential reason for the lack of auditory response is the noise generated by imaging gradients, which could have attenuated the auditory activity. Although measures were taken to ensure that the auditory volume was loud enough for participants to hear the sounds despite the scanner noise, the overall level of gradient noise may have led to a saturation of auditory activity, preventing us from detecting more subtle differences between the auditory stimulation and its absence. In an additional control analysis, we confirmed that our auditory stimuli evoked activity in the auditory cortex. However, the corresponding effects in the claustrum could have been smaller and thus harder to detect. Future studies aiming at measuring reliable auditory activity within the human claustrum could therefore take advantage of dedicated quiet EPI acquisition techniques that are tailored for fMRI studies of auditory processing ^32,33^.

Finally, it is possible that the temporal pattern of auditory responses in the claustrum neurons is different from that of the auditory cortex. A single-cell physiology study investigating claustrum responses to natural vocalizations observed that claustrum responses were highest when the vocalisations occurred immediately following silence, which may point to the claustrum’s role in change detection, an aspect closely related to attention ^26^. Accordingly, the claustrum may only show a response in the auditory modality when there is a salient change from one stimulus to the next. In our experiment, the auditory stimuli, which consisted of natural sounds (e.g., waterfalls, cat vocalisations, cowbells), may have not been sufficiently salient and behaviorally relevant to induce a response within the auditory claustrum. Future studies that focus on auditory processing within the claustrum could test the role of saliency in evoking stronger auditory responses.

Surprisingly, we found a deactivation in response to auditory stimuli in the right claustrum. This effect was observed in the right hemisphere only and was less consistent between sessions compared to visual activations. It should therefore be interpreted with caution and followed up in future studies. At this point we can only speculate about the potential functional significance of these deactivations. One possibility is that deactivation in response to auditory stimuli mirrors the well-known multisensory effects in the primary sensory cortices. It has been repeatedly shown that the presentation of stimuli in one modality leads to a deactivation in primary sensory cortex of the other modality ^34,35^. In this scenario, we would expect visual activations and auditory deactivations to coincide spatially, which is not the case (**Fig. 3c**). In addition, multisensory effects in the sensory cortices alone do not explain why deactivation only occurred in the right claustrum but not in the left one.

Another potential explanation is that these findings are due to attention-related processing within the claustrum and reflects distractor suppression, similar to what has been described in rodent literature ^13,14^. In our case, the central fixation task was used primarily as a way to ensure participants were fixating their gaze at the center of the screen. However, it is possible that because participants had to pay attention to the task, and hence to the visual modality, auditory stimulation served as a distractor stimulus, leading to suppression of the corresponding auditory representation within the claustrum whenever auditory stimulation occurred. This idea is consistent with fMRI findings in humans, which linked claustrum to within-modal and cross-modal divided attention ^36^ and task control ^7^ using conventional fMRI.

Our study did not yield any evidence for multisensory responses within the claustrum. A previous study in primates also did not find any evidence of multisensory responses, at least in the visual and auditory claustrum zones ^9^. Single-cell recordings as used in that study are a unique possibility to measure activity of individual neurons with an unprecedented spatial and temporal resolution, but at the same time they limit the spatial extent of brain tissue that can be examined within the same individual. Given the topography of the claustrocortical connections, multisensory responses could reside outside of the sensory zones, for example in areas that project to or receive inputs from the classical multisensory cortical regions such as the temporo-parietal junction (TPJ) and the intraparietal sulcus (IPS) ^37–40^. High-resolution functional MRI allowed us to measure activity throughout the whole claustrum, yet we did not find any evidence for multisensory effects, even beyond the visual zone. As with the lack of auditory activity, we cannot entirely rule out that some subregion of the claustrum shows multisensory responses, but we were unable to detect them due to the limitations of our method. In future high-resolution studies, a detailed map of the claustrum’s connection topography could be established to constrain the search of multisensory responses to a more specific location within the claustrum.

Here we present the first study that has investigated the fine-scale functional response of the human claustrum using fMRI with high spatial resolution at ultra-high magnetic field. We demonstrate that it is possible to detect evoked visual activity within the human claustrum. Although further studies are needed to determine the functional role of auditory deactivations, our current results for the visual modality open the possibility of studying the claustrum’s contribution to visual processing.

## Methods

The study’s design and methods were preregistered prior to conducting the experiment on AsPredicted.org (https://aspredicted.org/hj4sr.pdf). Code and data are available on the Open Science Framework (https://osf.io/7ebm2/)

### Participants

Sixteen healthy participants were recruited to take part in this study. The exact number of participants was determined during preregistration and was informed by previous investigations utilizing 7 Tesla fMRI that focused on subcortical structures and visual perception with a sample size of 6-8 participants ^41,42^. One participant had to be excluded due to poor data quality in the functional scans. Therefore, we had a total sample size of 15 (mean age = 24.60 years, SD = 3.33 years, 11 females and 4 males, 14 right-handed). Participants had normal or corrected-to-normal visual acuity, had no history of neurological impairments and were not taking any medication at the time of participation. All participants gave written informed consent prior to participation. The study was approved by the ethics committee of the Medical University of Vienna and was conducted in accordance with the Declaration of Helsinki. Participants received monetary reimbursement for their participation.

### Stimuli and task

Stimuli were presented using PsychoPy v2021.2.3 ^43^ software on a MacBook pro 13" running macOS 12 Monterey. Visual stimulation was projected on an MRI-compatible rear projection screen using an XGA VPL FX40 projector (Sony Group Corporation, Minato, Tokyo, Japan). The display was situated inside the scanner bore, which participants viewed through a mirror attached to the head coil at a 45° angle. The display-to-mirror distance was about 148 cm. Video stimuli were displayed at the full size of the MRI-compatible display (46.5 × 37cm) at a 16:9 aspect ratio and subtended 17.5° × 14° visual angle with a fixation point situated in the center of the screen measuring 4 mm in diameter with a visual angle of 0.15°. The central fixation point changed color between red and 9 different shades of green every 500 ms in a pseudorandom fashion in which we removed consecutive duplicate colors, meaning that a new color would be presented every 500 ms. Participant’s task was to fixate on the central fixation point and to respond when the fixation point changed to the color red. Auditory stimulation was delivered using S15 MR compatible in-ear earphones (Sensimetrics Corporation, Woburn, MA, USA). Prior to the beginning of the first functional run, a soundcheck was carried out to ensure that a sufficient level of loudness was achieved despite the ongoing scanner noise during image acquisition. This required fMRI dummy scanning while participants listened to some audio and provided feedback to increase or decrease the volume accordingly.

Stimuli consisted of naturalistic video scenes with a superimposed central fixation point. The videos were selected from the website Pexels (https://www.pexels.com/), a digital media sharing website. All stimuli materials used in this experiment were published to Pexels under a free-to-use and free-to-modify license (CC0, creative commons zero license). We used 48 video stimuli of different species of animals or natural scenery, and we ensured that the videos contained motion such that they would not be perceived as still images (for an example of the stimuli (see https://osf.io/7ebm2/)). The original video clips were modified using FFmpeg version 2022-07-18-git-cb22d5ea3c-full_build- www.gyan.dev ^44^ as follows. Each of the 48 selected videos were first cut down to a length of 15 seconds and then modified twice to create a visual only and an auditory only version. For both the audiovisual and auditory only conditions the audio was normalized, such that the maximum sound volume for each clip did not exceed a max value of 0 dB. These modifications yielded a total of 144 unique 15-second clips each belonging to one of the three stimulus types, corresponding to the three experimental conditions: audiovisual (AV), visual only (V) and auditory only (A).

To ensure consistency of our findings within individuals, each subject took part in two scanning sessions taking place on two separate days, with the mean time between session 1 and session 2 of 10.67 days (SD = 6.39 days). Each session contained 6 runs, and in each run 24 trials with stimulation were presented to participants. Each trial belonged to one of the 3 experimental conditions, with the order of conditions pseudorandomized and counterbalanced using a first-order counterbalanced condition sequence, to minimize trial history effects ^45^. On every 7^th^ trial a baseline (no stimulus) was presented, resulting in a total of 28 trials per run (see **Fig. 1a**, for an example of the presentation order). Each trial (including baseline) lasted 15 seconds. Each functional run lasted 420 seconds and participants took part in 6 functional runs per session for a total on-task scan time of 42 minutes. Two of the pilot sessions included in the analysis only involved a single session with 4 runs, and 1 pilot participant carried out 3 sessions resulting in 11 runs. The pilot sessions were not included in the session-to-session consistency analysis.

## MRI acquisition

### Functional MRI acquisition

MRI data were acquired using an ultra-high field 7 Tesla Siemens MAGNETOM scanner (Siemens Healthineers, Erlangen, Germany) using a 32-channel head coil (Nova Medical, Wilmington, MA, USA). Blood oxygen level dependent (BOLD) contrast was obtained by using the gradient-recalled echo-planar imaging (GE-EPI) sequence. We acquired 37 sagittal slices at 1.34 mm × 1.34 mm resolution (slice thickness = 0.8 mm; TR = 2000 ms; TE = 23 ms; FA = 62°, GRAPPA acceleration factor = 2). The parameters deviated slightly in one of the sessions of one pilot participant (TR=2500 ms, 47 sagittal slices). This resulted in a partial brain coverage of either the left or right claustrum, depending on the participant. The EPI slice orientation and anisotropic voxel size were chosen to minimize blurring and maximize resolution along the left-right dimension, where claustrum is thinnest ^46^. The right claustrum was scanned in 6 of the participants and the left claustrum was scanned in 9 of the participants (including 3 pilot participants). Each run of the functional scan took 420 seconds to complete, yielding 220 volumes per run. In addition to the main experimental runs, two additional volumes with identical parameters and opposite phase-encoding directions were acquired for subsequent susceptibility distortion correction.

### Anatomical MRI acquisition

Anatomical images were acquired using a T1-weighted MP2RAGE sequence ^47^ with 0.75 mm isotropic voxel size (matrix size: 320 × 300 slices: TR = 4300 ms; TE = 2.27 ms; FA = 4°, TI1 = 1000 ms, TI2 = 3200 ms, GRAPPA R = 2). The total time taken to complete the MP2RAGE scan was 530 seconds. Anatomical scans were obtained during both the first and second scanning sessions except for pilots with 1 session and for the pilot with 3 sessions (we used 2 of the anatomical scans for the latter).

## Data Analysis

### MRI data preprocessing

The preprocessing of the anatomical scans consisted of the following steps. Each of the two anatomical scans were first processed with the ‘presurfer’ tool (https://github.com/srikash/presurfer) to remove extracerebral noise from the MP2RAGE image by utilizing a bias-corrected second inversion of the MP2RAGE acquisition. After this, the two T1- weighted scans were co-registered using the robust registration method ^48^, which is a part of the FreeSurfer package, and averaged to produce one final structural image per subject. This image was passed to the CAT 12.8.1 toolbox ^49^ in SPM12 (7771, 13 Jan, 2020) (http://www.fil.ion.ucl.ac.uk/spm/, 2011) ^50^ running on MATLAB R2019b (MathWorks, Natick, MA, 2019). This was done to create a high-quality brain mask by concatenating the white matter (WM) and grey matter (GM) segmentations. Each subject’s structural image was then used to perform cortical surface reconstruction using FreeSurfer’s recon-all stream ^52,53^ at native resolution ^54^, substituting the FreeSurfer’s auto-generated brain mask with the CAT12-derived brain mask.

Preprocessing of functional data involved motion correction, distortion correction, co-registration with the structural image, resampling to the anatomical image and normalization to MNI space. The software used for each of the functional preprocessing steps is listed in Table 1 below.

**Table 2.**
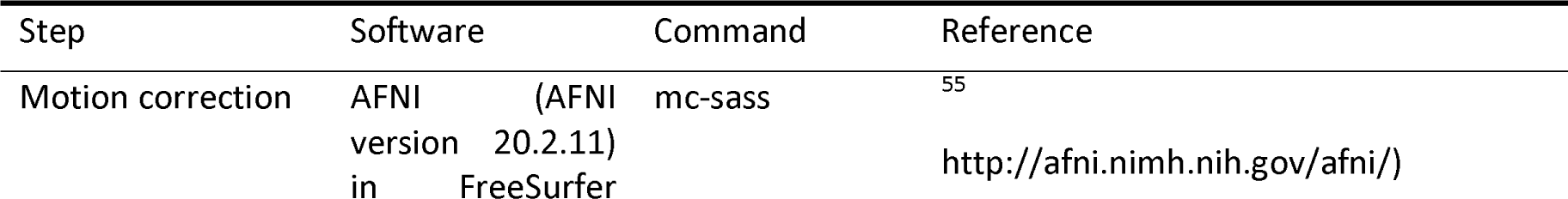
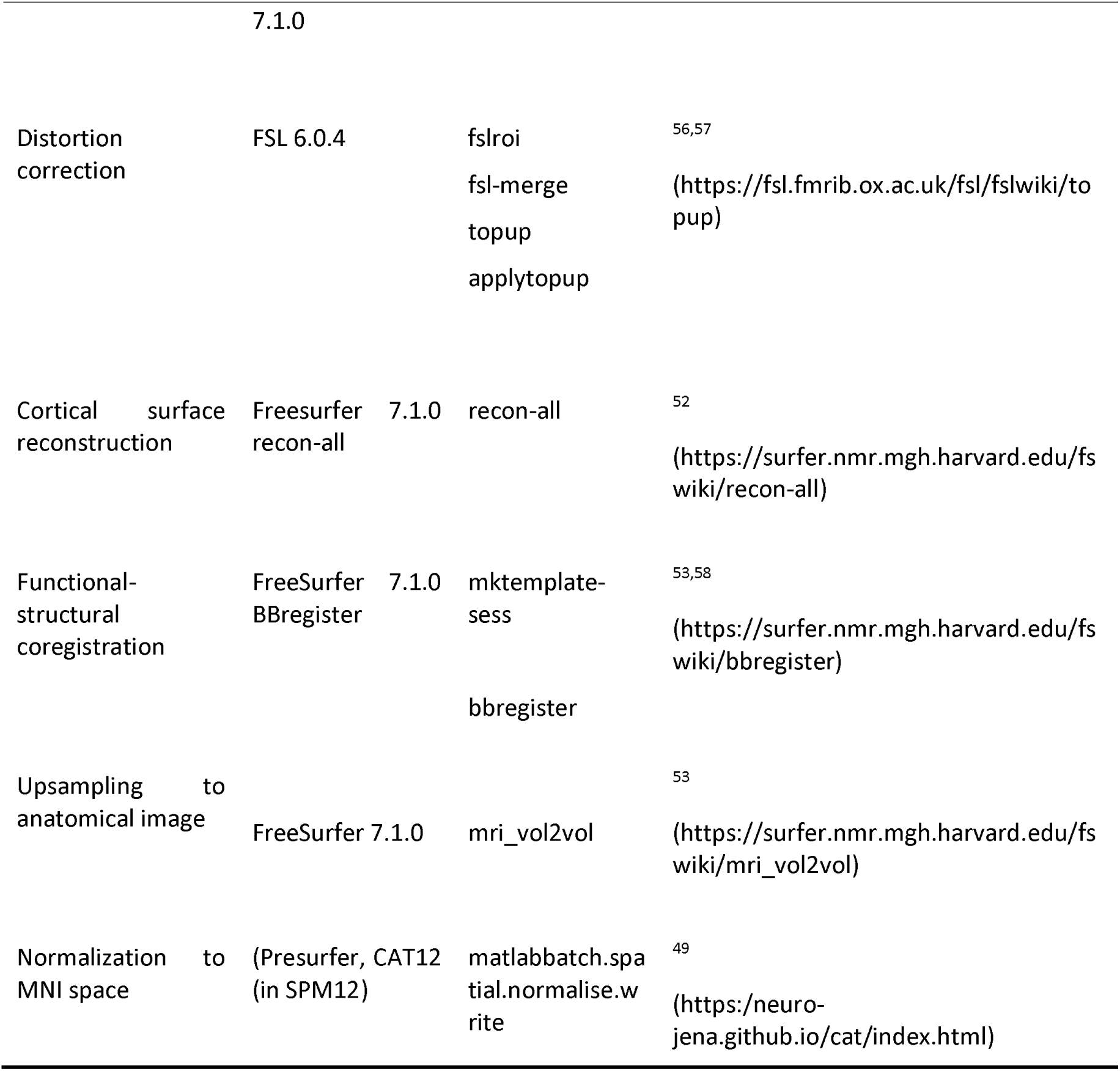
Steps involved in the preprocessing of functional data.

First, the images were corrected for subject motion using the AFNI 3dvolreg algorithm that is implemented as part of FreeSurfer’s functional analysis stream (FSFAST). After this, the functional images were corrected for susceptibility distortions due to magnetic field inhomogeneities using FSL’s ‘*topup’* and ‘*applytopup*’ ^56^. Distortion-corrected functional images were co-registered to the anatomical average scan using boundary-based registration with 6 degrees-of-freedom ^53^. We then resampled the functional data to the resolution of the structural scan (isotropic voxel size 0.75 mm) using FreeSurfer’s ‘*mri_vol2vol*’ command. This was done to bring the data from the two sessions into the same space in order to conduct the joint first-level analysis of both sessions. Normalization of the functional data to MNI space was performed by applying the nonlinear transformation derived from the CAT12 toolbox during anatomical processing. MNI-space data was additionally smoothed by convolving each volume with a 3D Gaussian kernel with a full width at half maximum (FWHM) of 1.5 mm using FreeSurfer’s ‘*mri_fwhm*’ command. Smoothed data were used for the group voxel-wise GLM analysis. Unsmoothed data were used for individual subject analysis (reporting individual peak activity of each subject and session-to-session consistency of voxel selectivity).

### Claustrum definition

To determine whether activations we observed in the functional experiment are located within the claustrum, it was necessary to accurately define the left and the right claustrum in each individual. Due to the fact that the standard atlases either do not contain a claustrum label at all ^59–61^ or focus exclusively on the dorsal claustrum part ^62,63^, we manually labelled the left and right claustrum in an ultra-high resolution (0.1 mm isotropic) 7T post-mortem brain dataset, which is available in MNI space ^21^. Labelling was performed using FreeSurfer version 7.1.0 FreeView tool. The photographs of Nissl-stained sections obtained from BrainMaps.org ^64^ and label schematics from the Atlas of the Human Brain ^65^ were used as a reference. Care was taken to ensure that the label included the ventral claustrum, an area which is difficult to differentiate using 3T and even 7T T1-weighted in vivo anatomical scans due to low gray matter density ^66–68^. Each claustrum was labelled twice by two independent research trainees. To quantify the consistency between the labelers, we calculated the dice similarity coefficient (DSC) between the two labels for the left and the right claustrum. For the left label the dice similarity coefficient between the two labels was DSC = 0.76 and for the right claustrum the dice similarity coefficient was DSC = 0.68 and the union between the two individual labels was used as the final claustrum ROI (**Fig. 1b**). After this, each label was smoothed by a convolution with Gaussian kernel of 0.1 mm FWHM using ‘*mri_fwhm*’. We then binarized the label using ‘*mri_binarize*’ in FreeSurfer and then performed the union using ‘*fsl_maths*’ command in FSL ^57^. The union of the left and right claustrum label was then resampled to the anatomical space for use as a mask for statistical inference in group analysis and as masks for individual subject plots (see results section).

To ensure that the claustrum label derived from the MNI space did not include any of the neighboring structures in individual subject space (e.g., insula, putamen), we projected the MNI-space label into each subject’s individual space by applying the inverse transformation generated by CAT12. FreeSurfer’s cortical-subcortical segmentation (aseg.mgz) was then used to visually check whether any of the claustrum voxels overlaps with structures other than voxels labelled as white matter (because the claustrum label is not available in FreeSurfer). We found that oftentimes the putamen label overlapped with the claustrum. However, a closer visual inspection showed that this was an issue with FreeSurfer automatic cortical-subcortical segmentation.

### Individual-level analysis

Individual-level analysis was performed using FreeSurfer’s FSFAST using a standard GLM approach with Visual (V), auditory (A), audiovisual (AV) and baseline conditions as regressors of interest, which were convolved with the canonical hemodynamic response function. In addition, run-specific offsets, scanner drifts (modelled with quadratic polynomial term) and the first 4 timepoints (to ensure that the scanner reached magnetic equilibrium) were modelled as nuisance regressors. To identify voxels within the claustrum that showed preference for either visual or auditory stimuli, we compared the beta estimates for the corresponding regressors, yielding the following contrasts: V − baseline and A

− baseline. To identify voxels responding to both modalities, we also calculated a multisensory contrast. Following Noppeney ^23^, we looked for a super-additive effect of the multisensory condition by calculating the contrast (AV + baseline) − (A + V). A GLM fit and contrast calculation was first performed individually for each session. The result of each session was then combined by carrying out a subject-level fixed-effects GLM analysis for each contrast.

### Group analysis

For the second-level group analysis, we used the contrast estimates from the individual-level GLMs to perform one-sample t-tests for nonzero effects using the random-effects GLM. Since the visual and auditory conditions are expected to activate a small part of the whole claustrum, we corrected the results for multiple comparisons within the claustrum label using cluster-wise permutations ^69^. We carried out 1000 permutations using a cluster-forming threshold of p<0.01 and a cluster significance level of p<0.05 ^70^ for each contrast. Since the left and the right claustrum data cannot be combined in this approach, group analysis was performed twice, once for subjects with the left claustrum scans (n = 9) and once for subjects with the right claustrum scans (n = 6).

### Session-to-session consistency analysis

In order to determine whether the visual and auditory responses within the claustrum of each individual appear consistently at the same location across both sessions 1 and session 2, we performed an analysis in which we used data from either session to define a region of interest (ROI) and data from the contrary session to measure the responses in that ROI. For example, to measure the visual response in session 1 we used session 2 to define the visually responsive voxels (contrast “V − baseline”, p<0.05 uncorrected) and then extracted the average contrast estimates from these voxels in session 1. This procedure was repeated with session 2 by defining visually responsive voxels using data from session 1. The same analysis was performed for auditory responses. The whole procedure yielded 4 values for each of the 12 subjects: visual activations in session 1 and 2 and auditory activations in session 1 and 2. Additionally, we calculated session-to-session consistency for auditory deactivations in session 1 and 2 for the 6 subjects with only the right claustrum scanned. Subjects with either their left or right claustrum scanned were pooled together. Session-to-session consistency was assessed by testing the ROI responses against zero using a one-sample t-test.

### Control analysis of auditory cortex activity

Since we did not find auditory activations within the claustrum, we wanted to ensure that the auditory stimuli used in the experiment were sufficient to activate the auditory cortex. To achieve this, we defined voxels corresponding to the transverse temporal gyrus (Heschl’s gyrus) in every participant from the automatic segmentation generated by the FreeSurfer recon-all stream, e.g. aparc+aseg.nii.gz ^52,60^ using the ‘*mri_extractlabel*’ command. Using the ROI, we extracted average beta estimates for each condition and compared the auditory condition with baseline.

### Behavioral analysis

To ensure that the participants maintained stable gaze fixation throughout the experiment we examined behavioral responses of each participant. To do this, we examined a time period between 0 and 2000 ms after the onset of each red color to determine whether a response was given. A response made in this time window was classified as a “hit”. We removed accidental duplicate responses and kept only the first response participants made. We then calculated the percentage of hits 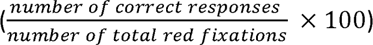 and mean reaction time for each participant. For two participants we encountered a technical issue with the button response box that failed to record responses during two runs of the first session. Therefore, for these two participants behavioral analysis was performed with the remining 4 unaffected runs only. Moreover, analysis of behavioral responses for the first pilot subject was not possible due to a technical issue with fixation color changes not being logged by the script. Since we did record button presses from this subject and the number of recorded presses was in the range of the remaining participants, it is unlikely that the participant did not perform the task properly. We thus report behavioral performance for 14 out of 15 subjects.

## Acknowledgements

The authors would like to thank Denis Chaimow for suggestions on MP2RAGE preprocessing, Joana Leitao for discussion of the multisensory integration analysis, as well as Erik Fink, Stefan Hödl, Maximilan Gerschütz and Elias Märzendorfer for their help in creating the claustrum label. This study was funded by the BioTechMed-Graz Young Researcher Group Grant to N.Z.

## Author information

## Authors and affiliations

**Institute of Psychology, University of Graz, Graz, Austria**

**BioTechMed, Graz, Austria**

Adam Coates, Anja Ischebeck & Natalia Zaretskaya

**High-Field MR Center, Center for Medical Physics and Biomedical Engineering, Medical University of Vienna, Vienna, Austria**

David Linhardt & Christian Windischberger

## Contributions

A.C. contributed to conceptualization, data analysis, original draft writing, reviewing and editing and methodology. D.L. contributed to data collection, review and editing and methodology. C.W. contributed to review and editing and methodology. A.I. contributed to supervision, review and editing. N.Z. contributed to conceptualization, funding resources, project administration, resources, methodology, supervision, review and editing.

### Corresponding authors

Adam Coates, Institute of Psychology, University of Graz, Universitätsplatz 2, 8010 Graz, Austria.

Natalia Zaretskaya, Institute of Psychology, University of Graz, Universitätsplatz 2, 8010 Graz, Austria.

### Ethics declarations

