## Supplementary Figures for "High-resolution 7T fMRI reveals the visual sensory zone of the human claustrum"

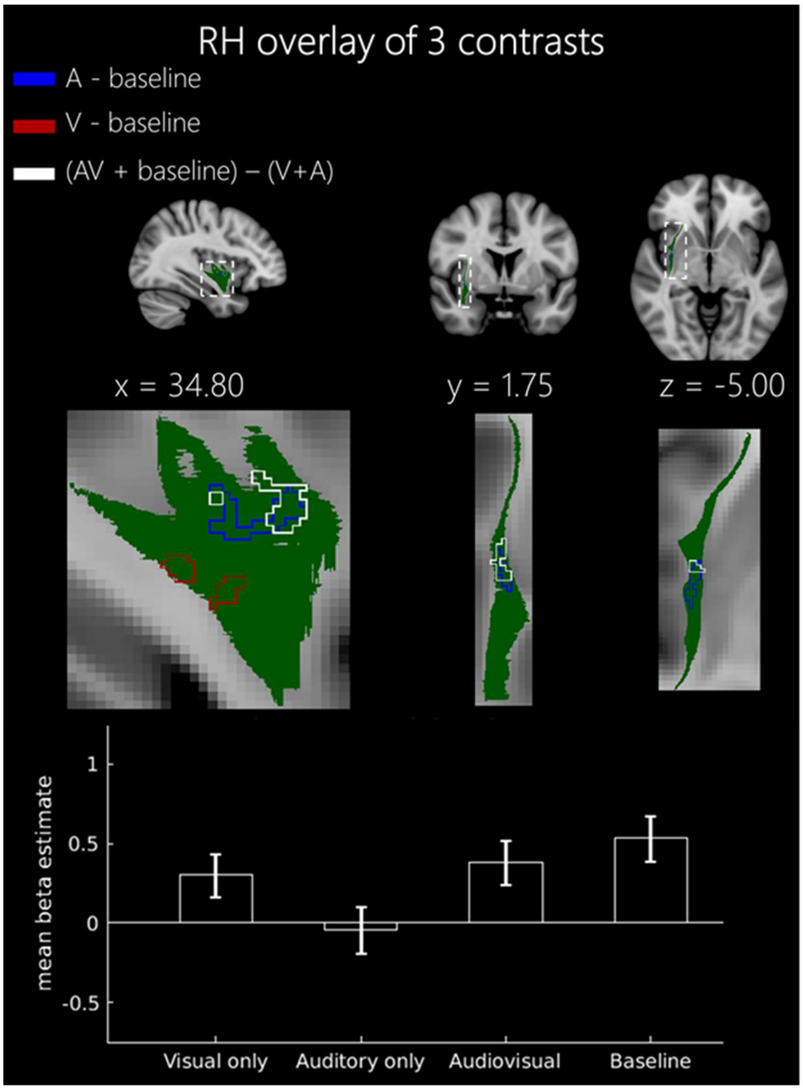


*Fig 1. Right claustrum auditory deactivations (blue), visual activations (red) and multisensory contrasts (white) overlayed onto the MNI template are shown as outlines to demonstrate the location and overlap of these effects. Bar plot represents the mean beta estimates for each experimental condition within the significant voxels of the right claustrum for the superadditive contrast (AV + baseline) – (V + A). Error bars represent SEM.*
